# Utilizing Non-Invasive Prenatal Test Sequencing Data Resource for Human Genetic Investigation

**DOI:** 10.1101/2023.12.11.570976

**Authors:** Siyang Liu, Shujia Huang, Yanhong Liu, Yuqin Gu, Xingchen Lin, Huanhuan Zhu, Hankui Liu, Zhe Xu, Shiyao Cheng, Xianmei Lan, Linxuan Li, Guo-Bo Chen, Hao Li, Xun Xu, Rasmus Nielsen, Robert W Davies, Anders Albrechtsen, Xiu Qiu, Xin Jin

**Affiliations:** School of Public Health (Shenzhen), Shenzhen Campus of Sun Yat-sen University, Shenzhen, 518107, China; BGI-Shenzhen, Shenzhen 518083, Guangdong, China; Division of Birth Cohort Study, Guangzhou Women and Children’s Medical Center, Guangzhou Medical University, Guangzhou, 510623, China; BGI Genomics, BGI-Shenzhen, Shenzhen 518083, Guangdong, China; Department of Neurology, Beijing Tiantan Hospital, Capital Medical University, Beijing 100070, China; College of Life Sciences, University of Chinese Academy of Sciences, Beijing 100049, China; Center for Productive Medicine, Department of Genetic and Genomic Medicine, Clinical Research Institute, Zhejiang Provincial People’s Hospital, People’s Hospital of Hangzhou Medical College, Hangzhou 310014, Zhejiang, China; Department of Integrative Biology, University of California, Berkeley, Berkeley, CA 94720, USA; Department of Statistics, University of Oxford, Oxford, United Kingdom; Bioinformatics Centre, Department of Biology, University of Copenhagen, Copenhagen 2200, Denmark; Provincial Clinical Research Center for Child Health, Guangzhou, 510623, China; Department of Women’s Health, Provincial Key Clinical Specialty of Woman and Child Health, Guangzhou Women and Children’s Medical Center, Guangzhou Medical University, Guangzhou, 510623, China; School of Medicine, South China University of Technology, Guangzhou 510006, Guangdong, China

**Keywords:** Non-invasive prenatal test sequencing, variant detection, genotype imputation, genome-wide association analysis

## Abstract

Non-invasive prenatal testing (NIPT) employs ultra-low-pass sequencing of maternal plasma cell-free DNA to detect fetal trisomy. With exceptional sensitivity, specificity, and safety, NIPT has gained global adoption, exceeding ten million tests, establishing it as one of the largest human genetic resources. This resource holds immense potential for exploring population genetic variations and their correlations with phenotypes. Here, we present comprehensive methods tailored for analyzing large, low-depth NIPT genetic datasets, involving customized algorithms and software for genetic variation detection, genotype imputation, and genome-wide association analysis. Through evaluations, we demonstrate that, when integrated with appropriate probabilistic models and population-specific haplotype reference panels, accurate allele frequency estimation and high genotype imputation accuracy (0.8 to 0.9) are achievable for genetic variants with alternative allele frequencies between 0.01 and 0.05, at sequencing depths of 0.1x to 0.25x. Additionally, we attained an R-square exceeding 0.9 for estimating genetic effect sizes across various sequencing platforms. These findings establish a robust theoretical and practical foundation for leveraging NIPT data in advancing medical genetic studies, not only in realms of maternal and child health, but also for long-term health outcomes.

**Highlights:**

- Introduction of probabilistic model integration for analyzing large-scale, low-pass non-invasive prenatal test (NIPT) sequencing data
- Evaluation of protocols for variant detection, genotype imputation, and genome-wide association analyses with NIPT data

## Introduction

Genetic variation plays a pivotal role in determining individual susceptibility to traits and diseases, with the intricate relationship between sequence variation and disease predisposition serving as a potent tool for understanding disease pathogenesis and developing innovative approaches to prevention and treatment. However, progress in genomics and multi-omics studies is hindered, particularly in low- and middle-income countries, by logistical and financial constraints that impede representative sampling from the entire population^1^. The predominant focus on individuals of European descent has resulted in a gap in explaining extensive trait variability and disease susceptibility across diverse populations^2^. Additionally, the predominantly cross-sectional nature of studies limits our understanding of the variability of genetic effects throughout life, subject to modification by aging, environments, and critical periods such as pregnancy^3^.

In recent years, non-invasive prenatal testing (NIPT) sequencing has ushered in a paradigm shift in pregnancy screening programs, offering a safer and more accurate alternative to conventional invasive procedures^4,5^. This transformative technology has been globally integrated into pregnancy screening programs, resulting in an unprecedented accumulation of genetic data that stands as a significant global genetic resource^6^. The core principle of NIPT involves the non-invasive acquisition of genetic information from peripheral blood samples from pregnant women, commencing as early as the 12th gestational week (**Figure 1a**). Several key characteristics define NIPT data. Firstly, an average of 89.99% of the DNA originates from mothers, while 10.01% is fetal DNA (**Figure S1**). Therefore, although NIPT was initially developed to detect fetal trisomy, the genetic information it contains predominantly reflects the maternal genome. Secondly, the sequencing depth generally ranges from 0.06x to 0.5x in common clinical settings. In China, data from several hospitals indicate that the average sequencing depth from Illumina supplier is 0.06 fold, while depth from BGISEQ and Ion Torrent suppliers ranged from 0.15 fold to 0.27 fold on average (**Figure 1b**). Thirdly, given the intensive 40-week examination period following mandatory pregnancy screening programs in various countries^7^, DNA information is correlated with enriched pregnancy phenotypes (**Figure 1c**). These phenotypes encompass standard physical examinations like height, weight and blood pressure, a typical set of approximately a hundred biomarkers widely utilized in pregnancy screening programs, such as glucose and lipid levels, recorded disease conditions in electronic medical records, and multi-omics data like metabolomes of mothers and children. The matching of genotypes and phenotypes in vast datasets significantly enhances the utility of NIPT data for investigating the genetic basis of pregnancy disorders, estimating genetic effects, and establishing polygenic risk scores for common biomarkers. As the volume of NIPT data continues to escalate, it becomes increasingly evident that realizing its full potential requires a systematic and rigorous approach to data analysis. Despite its profound significance, there exists a conspicuous void characterized by the absence of a comprehensive exposition and publication of methodologies tailored specifically for the analysis of NIPT data in human genetics investigations.

**Figure 1.**
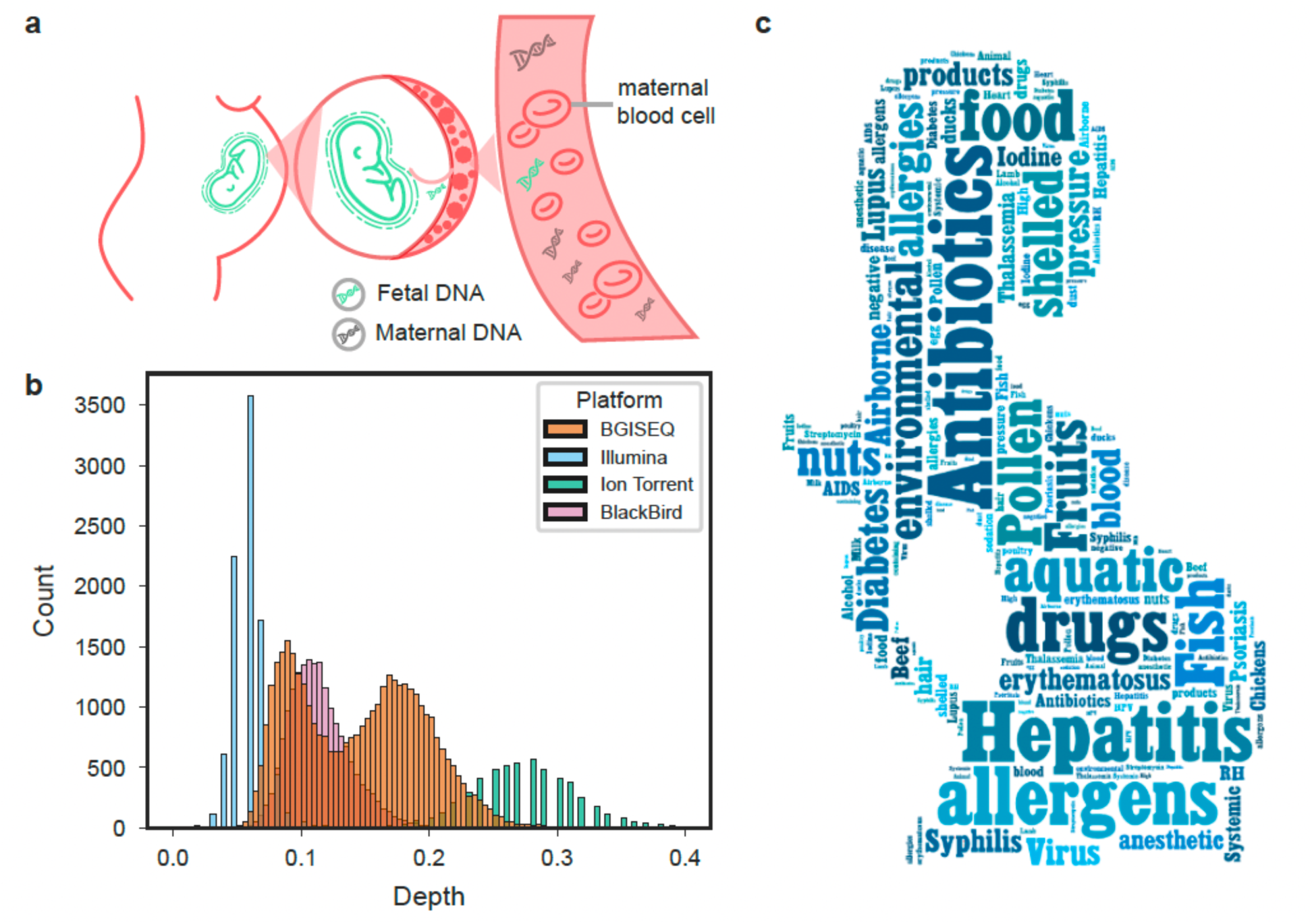
Characteristics of Standard Non-Invasive Prenatal Test Sequencing Data. (a) NIPT sequencing entails the sequencing and analysis of peripheral blood samples obtained from pregnant women. (b) Visualization depicting the typical sequencing depth observed in clinical settings for various sequencing platforms commonly used in China. (c) Integration of non-invasive prenatal test sequencing into pregnancy screening programs in China and other countries has enabled a connection with diverse maternal and children’s phenotypes.

In response to this critical gap, our study is dedicated to addressing the methodological challenges associated with NIPT data analysis head-on. We present a meticulous and systematic evaluation of various analytical methods, encompassing genetic detection, genotype imputation, the assessment of family relatedness, principal component analysis, and genome-wide association studies. Recognizing the imperative nature of resource sharing and collaboration in scientific endeavors, we are releasing our analytical pipeline, accompanied by a comprehensive protocol, to facilitate the analysis of NIPT data. Through these efforts, our objective is to provide researchers and scientists worldwide with a robust framework that empowers them to derive meaningful insights from NIPT data.

## Results

### Maximum likelihood model for SNP discovery and allele frequency estimation with NIPT data

The traditional multi-sample variation calling algorithms (samtools-bcftools^8^ and gatk unifiedgenotyper^9^) typically concentrate on bi-allelic alleles. These algorithms employ Bayesian approaches for variant discovery and genotyping, simultaneously estimating the probability that the two alleles- the reference allele and the alternative allele- are segregating in a sample of N individuals, and the likelihoods for each of the AA, AB, BB genotypes for each individual. In scenarios involving multiple-allelic probability estimations, each individual can have a maximum of ten potential combinations of genotypes. However, for very low-coverage sequencing, such as 0.1x, genotype likelihood may be unnecessary without associated haplotype information. Therefore, instead of estimating the likelihood of ten possible genotype combinations, we simplify the complexity by estimating the likelihood of four possible bases for the sampled one read from each individual. For a specific locus, the overall marginal data likelihood of N individual given the observed bases, base quality, and estimated population allele frequency can be aggregated by individual data likelihood, which is the sum of the probability of the four possible bases (**Figure S2** and Methods). The variant detection algorithm has been implemented in BaseVar (see Code Availability). In a simulation study, we demonstrate that BaseVar can robustly identify variants given a specific allele frequency threshold (**Figure 2**). The call rate and accuracy depend on the sample size, true alternative allele frequency, and the allelic type of the variant (bi-allelic, tri-allelic and tetra-allelic). For sample sizes of 44K, 140K and 1 million, we detected 100% of the bi-allelic variants with minimum alternative allele frequency of 0.015, 0.006 and 0.003, respectively (**Figure 2a and Table S2**). For a fixed sample size of 140K, we identified 100% of the bi-allelic, tri-allelic and tetra-allelic variants with minimum alternative allele frequencies of 0.006, 0.008 and 0.008, respectively (**Figure 2b**). The accuracy of alternative allele frequency estimation is high for all scenarios, with root mean square deviation (RMSD) ranging from 0.001 to 0.007 and normalized RMSD ranging from 0 to 1.3 (**Figure 2c-d, Table S2-S3**). In a comparison of the accuracy of variant detection and allele frequency estimation for 2,504 individuals from the 1KG project^10^ among the BaseVar algorithms, Unifiedgenotyper, and samtools-bcftools, all three algorithms provide accurate variant detection and allele frequency estimation, with BaseVar slightly outperforming the other two softwares (**Figure S3**). Notably, when the sample size exceeds 100,000, UnifiedGenotyper and Samtools are not able to produce outcomes and we can only provide computation performance evaluation of BaseVar (**Table S4**).

**Figure 2.**
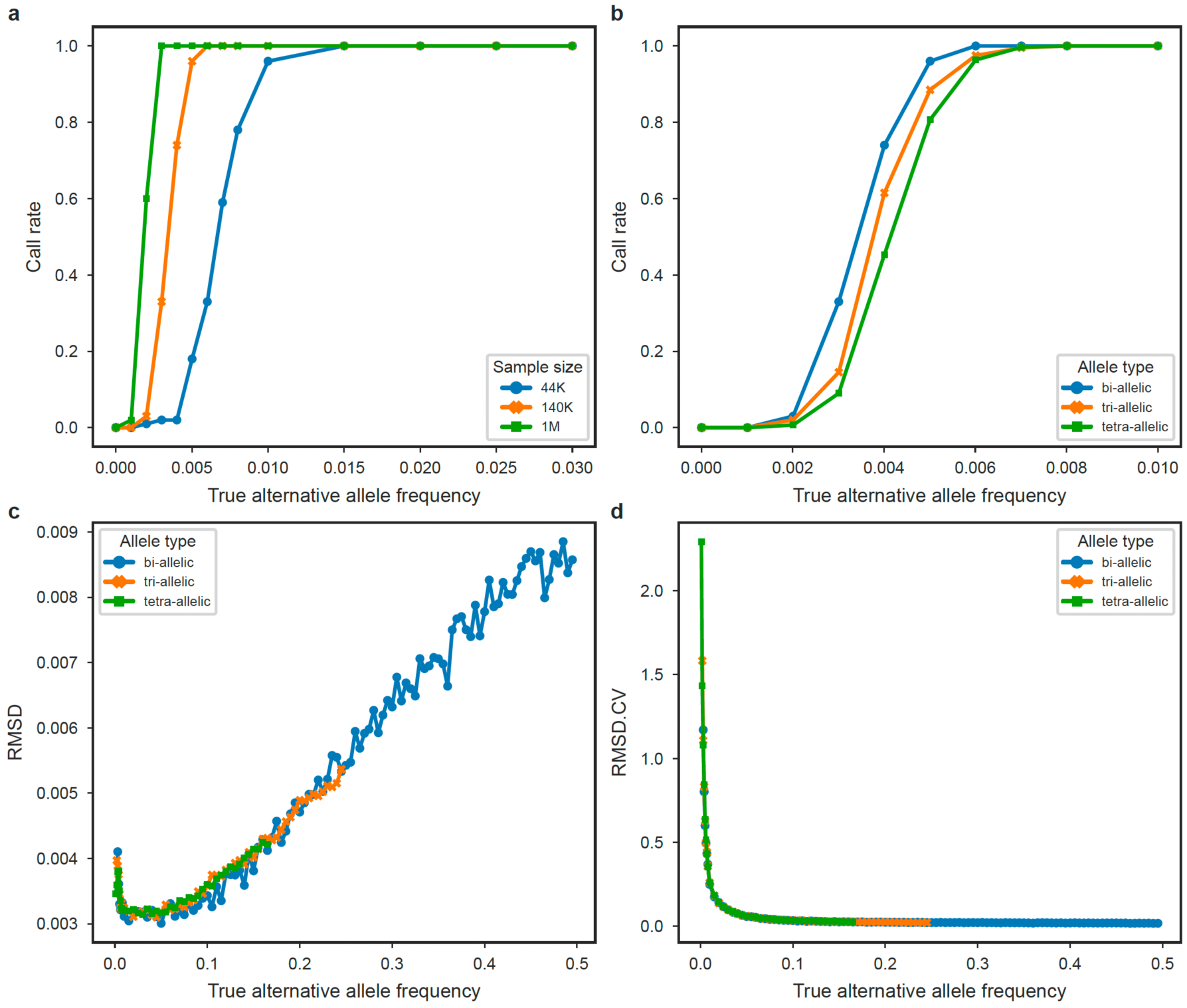
Assessment of Call Rate and Allele Frequency Accuracy in Relation to Allele Frequency on Simulation Data. (a) Call rates for three datasets, each comprising 44 thousand, 140 thousand, and 1 million individuals, focusing on bi-allelic variants. (b) Call rates for 140 thousand individuals across three allele types. (c) Root mean square deviation for allele frequency estimation. (d) Coefficient of variation for allele frequency estimation.

### Gibbs sampling and hidden markov model for genotype imputation

To leverage NIPT data for investigating the genetic architecture of maternal and child phenotypes, genotype imputation analysis is indispensable. QUILT^11^ and GLIMPSE^12^ present two robust algorithms specifically designed for low-pass whole genome sequencing data, accommodating the genotype uncertainty inherent in non-invasive prenatal sequencing data. Both methods employ a Hidden Markov Model with Gibbs sampling, utilizing prior allele frequency information derived from a haplotype reference panel. An essential question arises on what factors optimize genotype imputation accuracy? We conducted an evaluation of these two algorithms using three Chinese reference panels, using 110 NIPT samples, where whole blood was sequenced up to an average of 40x (the true set). These three reference panels consist of the 1000 Genome Project (1KGP) reference panel (N=504 unrelated East Asians, including 301 Chinese)^10^, the Born in Guangzhou cohort study (BIGCS) reference panel (N=2, Chinese with high-quality long-range phased haplotypes from duo and trio information)^13^ and the G10K reference panel (N=10,241 unrelated Chinese)^14^.

As depicted in **Figure 3**, imputation accuracy depends on the sample size in the reference panels and the average depth of the NIPT samples. Focusing on well-imputed variants (INFO score > 0.4) with the 1KGP reference panel, the QUILT algorithm achieved average genotype imputation accuracy of 0.80, 0.86 and 0.91 at average sequencing depths of 0.1x, 0.2x and 0.3x (**Figure 3, Table S5a**). GLIIMPSE performs slightly better than QUILT, exhibiting average genotype imputation accuracy of 0.82, 0.87 and 0.92 for sequencing depth of 0.1x, 0.2x and 0.3x, respectively (**Figure 3, Table S4b**). With an expansion of the reference sample size to thousands of related individuals (6x), genotype imputation accuracy improved. The average imputation accuracy increased to 0.84, 0.89 and 0.93 for NIPT sequencing depths of 0.1x, 0.2x and 0.3x with the GLIMPSE algorithm. Further improvement was observed when expanding the reference sample size to ten thousand individuals with high-coverage sequencing (∼40x), such as the STROMICS reference panel^14^. This demonstrated an average imputation accuracy increases to 0.84, 0.90 and 0.94 for NIPT sequencing depth of 0.1x, 0.2x and 0.3x with the Glimpse algorithm. Given the optimal performance with the STROMICS reference panel and assuming a million individuals are involved in a GWAS, a nine hundred thousand effective samples size can be achieved following genome-wide association power calculations (**Supplementary Notes**), which enables robust genome-wide association investigations (**Figure S4**).

**Figure 3.**
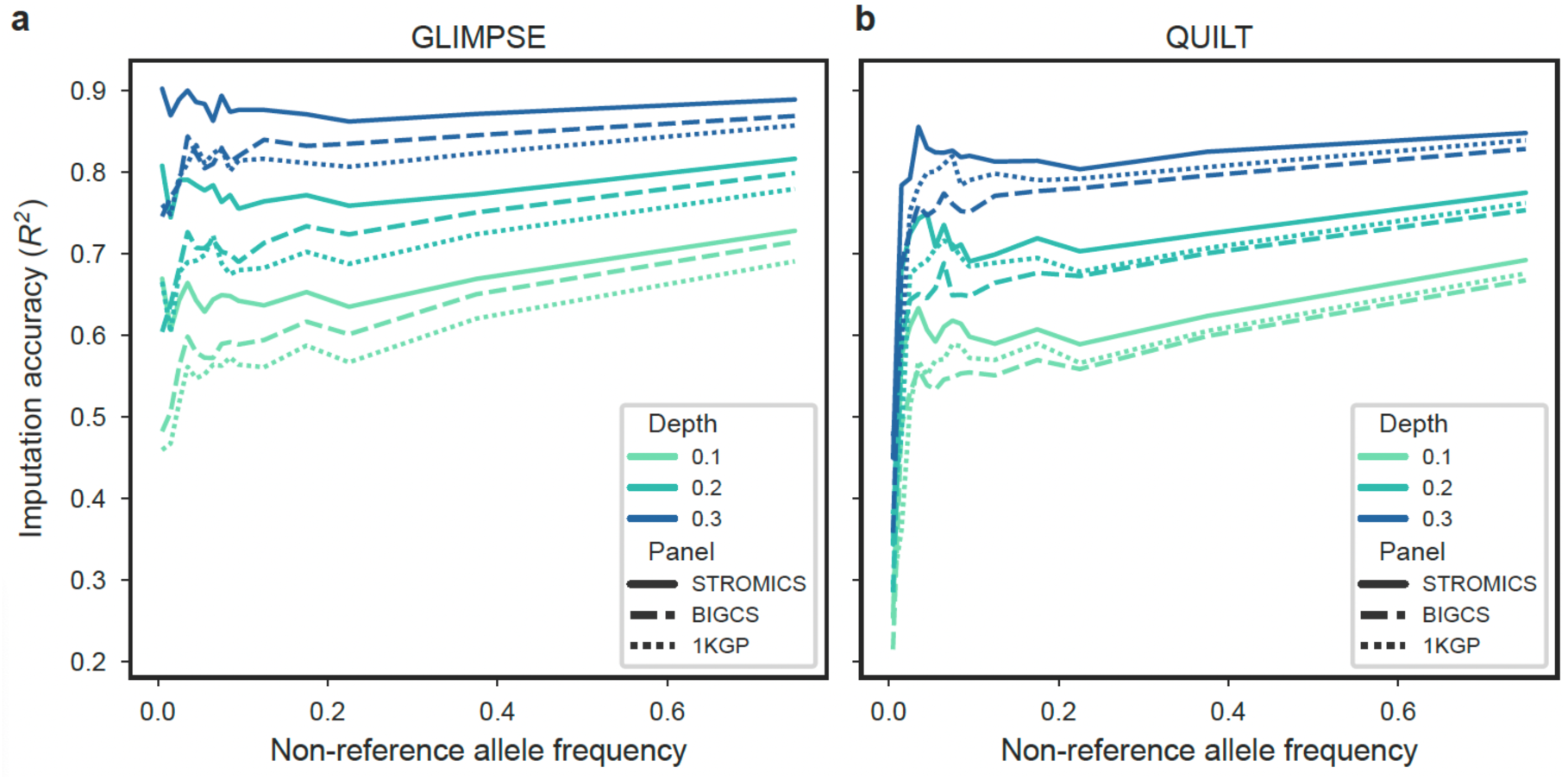
Imputation Accuracy of NIPT Samples Compared to High-Coverage True genomes. Imputation accuracy attained by the Glimpse algorithm (a) and the QUILT algorithm (b). The evaluation is performed against the reference panels of the 1000 Genomes Project, BIGCS, and STROMICS using bcftools. The typical NIPT sequencing depth spans from 0.1x to 0.3x.

### Family relatedness and population stratification

In genome-wide association studies, two crucial confounding factors are cryptic family relatedness and population stratification^15^. An important question pertains to the suitability of standard software, such as PLINK^16^ for examining family relatedness and population stratification. Additionally, it raises the issue of whether to utilize data before or after genotype imputation for analyzing the family relationships. To establish a protocol for investigating family relatedness investigation, we included 2,205 identical individuals, those tested more than once, to evaluate the kinship coefficient using data before and after genotype imputation. Results indicated an inability to generate correct estimates of the kinship coefficient with genotypes before imputation (**Figure 4a**). However, we were able to calculate correct kinship coefficient for genotypes after imputation. The kinship coefficients for all identical individuals exceeded 0.43, aligning with statistical expectations (the expectation of scaled kinship coefficient for duplicate samples is 0.5 in PLINK algorithm) (**Figure 4b**). Therefore, a convenient cutoff of ∼0.354 can be employed to screen for duplicate samples, and ∼0.177 can be used to filter first-degree relations in NIPT data, a scenario rarely encountered in NIPT data.

**Figure 4.**
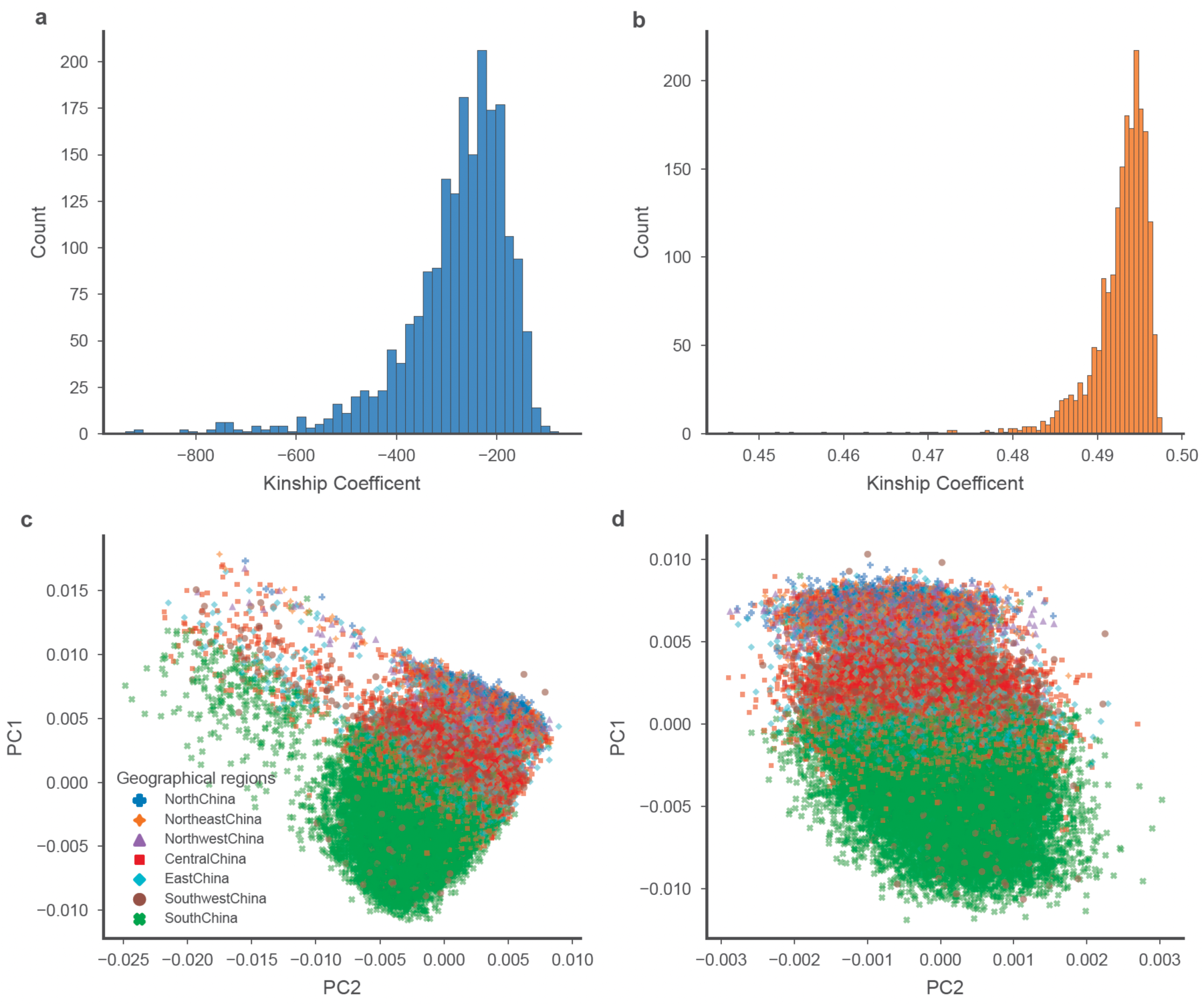
Family Relatedness and Population Stratification of NIPT Data. (a) Distribution of kinship coefficients for identical samples using PLINK with imputed genotypes (b) Distribution of kinship coefficients for identical samples using PLINK without imputed genotypes (c) Principal component analysis before genotype imputation (d) Principal component analysis after genotype imputation.

Concerning population stratification, we assessed principal component analysis (PCA) using the widely used PLINK algorithm and methods employing individual allele frequency (thus not relying on exact genotypes)^17^ for NIPT data. PCA was conducted with the PLINK algorithm, relying on exact genotypes before and after genotype imputation, as well as the EMU algorithm for a randomly selected 10,000 samples^18^. The first principal component analysis was observed to reflect latitudes across all scenarios (**Figure 4c-d, Figure S5**). Notably, both the EMU algorithm and the PLINK analysis on unimputed genotypes captured genotype missingness, that manifested as outliners. It is crucial to remove samples with excessively low sequencing depth and poor sequencing quality from the extensive pool of NIPT data.

### High consistency of genetic effect estimates among sequencing platforms

One of the most important applications of NIPT data lies in conducting Genome-wide association studies, particularly for traits readily accessible in maternal and child’s cohorts. The indispensable role of genotype imputation empowers robust GWAS. However, a remaining question pertains to the replicability and reliability of the significant loci identified in GWAS. To address this, we obtained GWAS results for the same maternal metabolite and newborn metabolite levels from four different sequencing platforms and compared the genetic effect of the significant loci (**Figure 5, Table S6**). Our findings reveal a remarkably high consistency in genome-wide association estimates between the BGI-seq500 and BlackBird ( 𝑟^!^ = 0.927 ) and between the Illumina sequencing and Ion Torren sequencing platforms (𝑟^!^ = 0.943). These statistics indicates a very high replicability of GWAS using the NIPT sequencing data.

**Figure 5.**
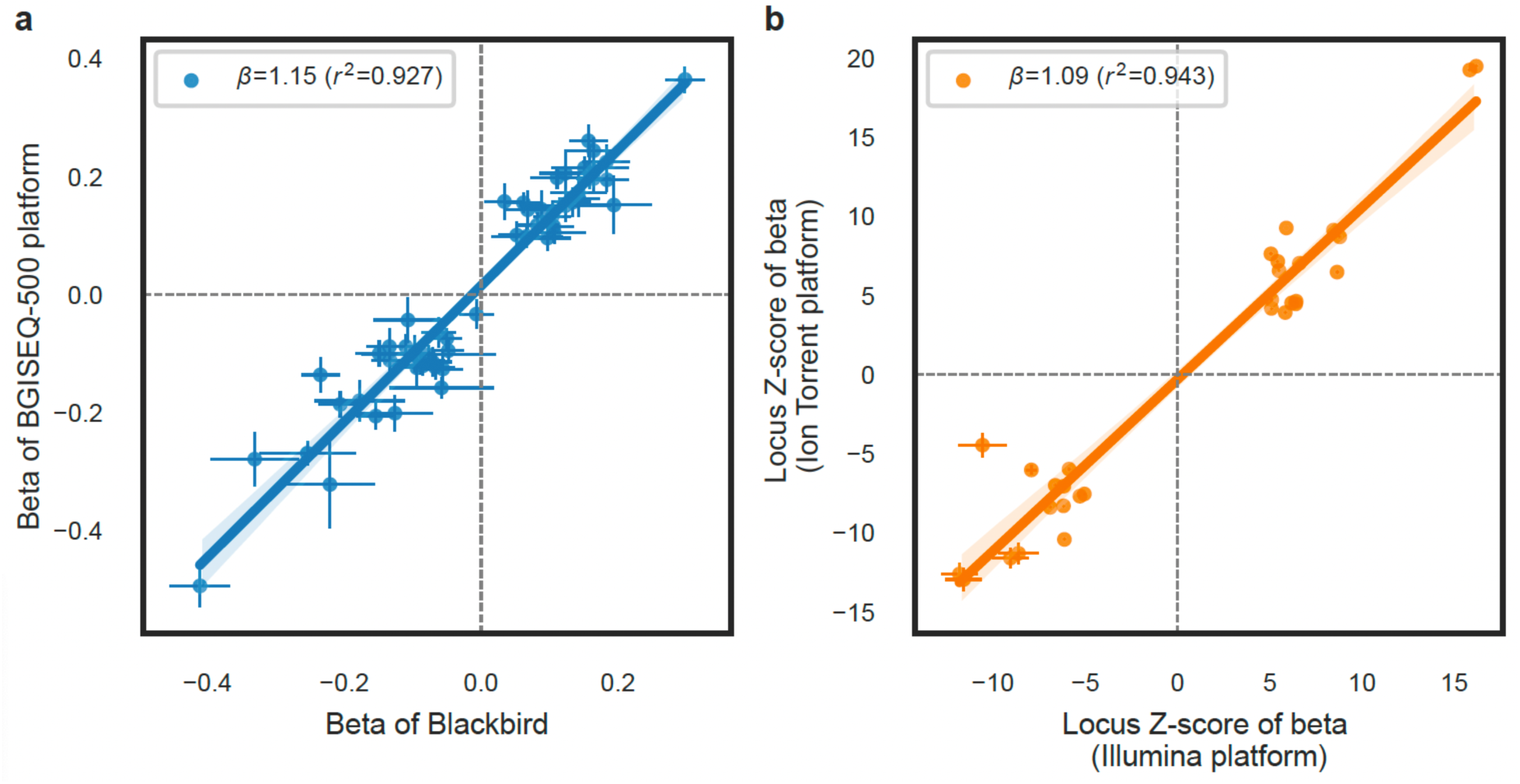
Consistency of genetic associations for different NIPT sequencing platforms. (a) Scatter plot showing the consistency of genetic effect obtain from genome-wide association studies between BGI-seq500 and BB platforms (b) Scatter plot showing the consistency of genetic effect between BGI-seq500 and the Z-score platforms.

## Discussion

NIPT data represents a new category of genomic data that is quickly mounted as the windfall of the historically fastest adopted molecular test-non-invasive prenatal testing (NIPT) that sequenced cell-free DNA of maternal plasma. Because of its fast accumulation at hospitals the sample size of a single NIPT cohort, typically around 50,000 samples, surpass those of chipped or sequenced cohorts. Despite the limitations of low-depth sequencing in driving individual discoveries, its potency for population and statistical genetic analyses is evident. This study presents a comprehensive suite of analytical methods tailored for human genetic investigations using extensive NIPT data. These methods encompass genetic variant detection, allele frequency estimation, genotype imputation, assessment of family relatedness, principal component analysis, and genome-wide association analyses.

Leveraging probabilistic modelling through maximum likelihood and likelihood ratio test algorithms, we achieve accurate detect genetic variant detection and precise estimation of allele frequencies. This facilitates in-depth exploration of site-specific allele frequencies in genomic databases^19^, enabling the derivation of regional allele frequencies and polygenic risk scores, which contributes to the discovery of varying genetic susceptibility among different populations^20^. Notably, our findings underscore the impact of NIPT sequencing depth and reference panel scale on genotype imputation performance, providing insights into achieving optimal accuracy. We observe that genotype imputation performance improves with increasing NIPT sequencing depth and the scale of the reference panel. The highest imputation accuracy achieved for the Chinese population is 0.84, 0.90 and 0.94 for NIPT data depths of 0.1, 0.2 and 0.3 fold, respectively, using a reference panel comprising over 10,000 individuals with high depth sequencing. Accurate inference of family relatedness for identical individuals is achieved with common software like PLINK, providing a uniform solution for NIPT data, where multiple pregnancy records for the same individuals are common, especially in large sample sizes. Principal component analyses on unimputed and imputed genotype matrices, as well as individual allele frequency matrices, reveal distinct data structure. PCA based on the unimputed genotype and individual allele frequency matrix captures both missingness levels and population genetic structure while PCA based on the imputed genotype matrix alleviates the missingness patterns. Importantly, we found that genome-wide association studies based on NIPT data from different sequencing platforms demonstrate highly consistent genetic effect estimation. The integrated methods are provided in the Code availability section.

These methodological advancements unlock the potential of NIPT data to address critical medical and biological questions. With the rapid accumulation and global prevalence of NIPT sequencing data, we anticipate its substantial contribution to diversifying populations in genetic studies. Specifically, NIPT data serves as a valuable resource for studying pregnancy-related phenotypes, forming the foundational basis for enhancing prenatal care and gaining deeper insights into common maternal and pediatric disorders. In several companion papers stemming from this investigation, we delve into the genetic associations with approximately a hundred biomarker phenotypes used in pregnancy screening^21^. Of notable importance is our exploration of the genetic basis underlying common pregnancy disorders, such as gestational diabetes^18,22^ and gestational thrombocytopenia^23^, particularly in the underrepresented Chinese population. Furthermore, we have garnered insights into the genetic architecture underpinning molecular phenotypes, including maternal^24^ and newborn metabolites^25^.

The publication of methods and shared experience arising from these practice lays the foundation and opens new avenues for future studies. In the clinical practice of NIPT testing, maternal blood plasma samples are stored in hospitals. Therefore, with informed consents, ethnic approval, and approval for collecting and utilizing genetic resources from specific authorities, such as the China Human Genetic Resources Administration Office in China, expanding the phenotypic collection becomes important. Beyond the readily available biomarkers and anthropometric parameters collected from pregnancy screening, there is potential to broaden the panel to include metabolites, lipidomes, and proteomes in the maternal plasma. Deciphering the genetic basis of the maternal phenome, when combined with birth cohort family sequencing data, enables investigations into maternal intrauterine and fetal genetic effects on birth outcome and, subsequently, long-term children’s health^26–28^. Secondly, it is worthwhile to enhance the sequencing depth and length of specific samples, especially those of case samples, to obtain a spectrum of Indel and SVs alongside SNP genetic markers underlying adverse pregnancy and birth outcomes^29,30^. Thirdly, for many women, pregnancy screening represents their first comprehensive physical examination in life. Therefore, combining this with long-term follow-up data allows for health predictions for participants. Fourthly, the increasing sequencing depth enables direct imputation of maternal and fetal genotypes from plasma sequencing samples, which is valuable for investigating the impact of fetal genotypes on the mothers and vice versa.

Throughout this process, the continuous development and evaluation of analytical methods and a suitable data sharing scheme specific to NIPT data ensure that researchers can effectively harness this resource, facilitating more precise and meaningful genetic research.

## Methods

### NIPT data collection

We acquired sequencing depth statistics from the following representative NIPT screening centers in China: BGI-Shenzhen Life Science Institute (utilizing BGIseq500 sequencing and Blackbird sequencing, N=39,194), Suzhou Maternal and Children’s Health hospital (employing Illumina sequencing, N = 8,960 and Ion Torrent sequencing, N=5,458), Wuhan Maternal and Children’s Health Hospital (BGIseq500 sequencing, N=39,178), Longgang District Maternity and Child Healthcare Hospital of Shenzhen City (BGIseq500 sequencing, N=70,739) and Shenzhen Baoan Women’s and Children’s Hospital (BGIseq500 sequencing, N=50,948) (Table S1).

The NIPT sequencing protocol can be briefly summarized as follows: Peripheral whole blood (5-10ml), approximately 5ug each, were drawn from each participant and stored in EDTA anticoagulant tubes to prevent hemolysis. Within 8 hours of blood collection, plasma was extracted from two rounds of centrifugation. The first round, conducted at 1,600g for 10 minutes, separated plasma from whole blood, and the second round, at 16,000g for 10 minutes, removed residual cells. Subsequently, plasma samples underwent library construction and sample quality assessment. Notably, cell-free DNA fragments were extracted from 0.6 ml plasma using the circulating nucleic acid kit (Qiagen, Germany). For Blackbird or BGI-seq500 sequencing platforms, a 36-cycle single-end multiplex sequencing approach was employed. For the Ion Proton platform, the sequencing library was constructed by an Ion plus fragment library kit (Life Technologies, USA), quantified with a qubit fluorometer, and sequenced in a 30-cycle run. Adapter sequences of reads were trimmed using the Ion Torrent platform-specific pipeline (Torrent Suite, version 2.0.1), generating reads of lengths ranging from 150 to 165 bp. Illumina platform sequencing involved library construction with a ChIP Seq library protocol, quantification with Kapa SYBR fast qPCR kit (Kapa Biosystems, Woburn, MA, USA), and single-end reads sequencing in a 37-cycle run on the Illumina HiSeq-2000 platform. Adapter sequences were trimmed, resulting in reads of 35 bp length. Quality control involved the removal of poor-quality reads using SOAPnuke (https://github.com/BGI-flexlab/SOAPnuke), with reads eliminated if they contained more than 30% low quality bases (Q<=2) or N bases. Overall, each participant underwent whole-genome sequencing yielding 5-10 million cleaned reads, corresponding to a sequencing depth of approximately 0.06x to 0.3x.

### High-coverage whole-genome sequencing of 100 participants

To facilitate an unbiased assessment of genotype imputation accuracy, we performed high-coverage whole-genome sequencing for an additional cohort of 100 healthy Chinese participants, which included the BGI-seq500 NIPT sequencing data. The sequencing was performed using the Illumina HiSeq X10 platform with 140bp paired-end reads, resulting in an average coverage of 40x. The obtained clean reads were aligned to the GRCh38/hg38 reference genome using BWA-MEM (v0.7.17)^31^. Subsequently, the GATK (v4.1.8.1) best practice joint calling protocol was applied to detect and genotype variants in these participants.

Following variant quality score recalibration (VQSR) and the removal of multi-allelic variants, we derived a set of 11,174,603 high-quality genotyped biallelic variants, consisting of 9,816,793 SNPs and 1,357,810 Indels. Further refinement involved excluding SNPs located within the low-complexity regions of GRCh38 and SNPs classified as singletons in these 100 participants. The resultant 8,303,052 SNP variants were used to access genotype imputation accuracy across the STROMICS, BIGCS and 1000 KGP reference panels.

### Variant detection

The performance of commonly used variant detection algorithms and software tools, including Unified Genotyper and Samtools, along with the BaseVar method for detecting single nucleotide polymorphisms and estimating allele frequency from low-pass sequencing data, was systematically assessed. This evaluation involved downsampling the genomic data of 2,504 individuals from the 1,000 Genomes Project to sequencing depths equivalent to those observed in NIPT data, ranging from 0.1x to 0.3x. The specific parameters employed for the three software tools are detailed below.

#### UnifiedGenotyper

The UnifiedGenotyper was executed with the following input parameters: “java -jar GenomeAnalysisTK.jar -T UnifiedGenotyper -R reference.fasta -nt 10 -I input.bam - stand_call_conf 30.0 -stand_emit_conf 0 -glm SNP -o output.vcf”.

#### Samtools-bcftools

Samtools was executed with the following input parameters: “bcftools mpileup -f reference.fa input.bam | bcftools call -mv -Ob -O z -o output.vcf.gz”.

#### BaseVar

The detailed algorithm of BaseVar is presented in Supplementary Notes. It was executed with the following input parameters: “basevar basetype -R reference.fasta --batch-count 50 -L bamfile.list --output-vcf test.vcf.gz --output-cvg test.cvg.tsv.gz --nCPU 4“ .

### Genotype imputation

#### QUILT and STITCH

We employed QUILT (version 1.0.4) to infer genotype probabilities from NIPT data. This process was conducted in 5-Mbp chunks with 250 kbp flanking buffers, specifically targeting the region on chromosome 20:1–64,444,167bp region. Hap and legend format files of reference haplotypes were generated from three haplotype VCFs (1KGP, BIGCS and STROMICS) using the bcftools “convert –haplegendsample” command. The CHS recombination rates file from 1KGP served as the genetic map file, with liftOver utilized to transition from chromosome position hg37 to hg38. Additional parameters, including “--nGibbsSamples=7, --n_seek_its=3, --nGen=1240, -- save_prepared_reference=TRUE” were set for the QUILT imputation. The initial values for the EM optimization of model parameters were based on allele frequency information from the 1KGP Chinese population (--reference_populations=CHB, CHS, CDX, N=301), the BIGCS reference panel (--reference_populations=BIGCS_PhaseI, N=2,243, constructed with family data) and the STROMICS reference panel (--reference_populations=STROMICS, N=10,241). Filtration thresholds were applied, requiring genotype quality greater than GQ15, depth (DP) greater than 10, IMPUTE2-style info score greater than 0.4, and a minor allele frequency greater than 0.001. It is noteworthy that QUILT and STITCH are two algorithms developed by the same author. While STITCH focuses on genotype imputation without a reference panel^11^, QUILT was specifically designed for genotype imputation with a reference panel^32^. Importantly, we would specifically note that starting from STITCH (version 1.2.7), the QULIT algorithm has been incorporated, representing a stable method before formally named QUILT.

#### GLIMPSE

For GLIMPSE (version 1.1.1), input data in the form of Genotype Likelihoods (GLs) was required. GLs were computed from sequencing data using BCFtools with the following commands: "bcftools mpileup -f ${REFGEN} -I -E -a ’FORMAT/DP’ -T ${reference_panel_VCF} -r chr22 ${BAM} -Ou" and "bcftools call -Aim -C alleles -T ${reference_panel_TSV} -Oz -o ${OUT}." These commands were applied to all target individuals and variant sites present in the reference panel of haplotypes used for imputation. Before initiating imputation and phasing, chunks for these processes were defined, with the minimal size of the imputation region was set at 2,000,000 base pairs, with a buffer region of at least 200,000 base pairs (GLIMPSE_chunk --input reference_panel

--region chr20 --window-size 2,000,000 --buffer-size 200,000 –output output_file). Subsequently, GLIMPSE_phase was run for each imputation chunk as separate jobs (GLIMPSE_phase –input ${VCF} --reference ${REF} --map ${MAP} --input-region ${IRG} --output-region ${ORG} -- output ${OUT}), utilizing genetic maps from "genetic_maps.b38" in GLIMPSE’s maps file. Finally, different chunks of the sample chromosome were merged using GLIMPSE_ligate.

### Family relatedness

PLINK2 (v2.00a3LM) was used to select SNPs with a minor allele frequency (MAF) of at least 5%. Kinship was calculated using PLINK2 based on the KING-robust kinship estimator with the following command: "plink2 --vcf vcf_file dosage=DS --maf 0.05 --make-king-table --out out_file." In the results, first-degree relations (parent-child, full siblings) correspond to approximately 0.25, second-degree relations correspond to about 0.125, and so forth. It is customary to use a cutoff of approximately 0.354 (the geometric mean of 0.5 and 0.25) to identify monozygotic twins and duplicate samples^16^.

### Population stratification

We applied PLINK2 (v2.00a3.6LM) for PCA both before and after genotype imputation. Additionally, EMU (v.0.9) was used for PCA specifically before genotype imputation, aiming to assess the genetic structure of a dataset comprising 70,608 samples from one of the sequencing centers^23^. All PCA analyses were conducted on a set of non-duplicated 3,843,382 biallelic SNP variants with MAF ≥ 5%. The PCA procedures were executed with the following commands for PLINK2 (plink2 --maf 0.05 --vcf vcf_file dosage=DS --pca 10 --remove duplicated.txt –out $outfile) and for EMU (EMU –mem –plink $infile --n_eig 10 –out $outfile).

### Evaluation of consistency in genetic effect estimates in genome-wide association analyses

We conducted a comparison of genetic effects for a combined total of 53 loci significantly associated with maternal metabolites^24^ and 30 loci significantly associated with neonate metabolites^25^. These loci were identified in independent sets of individuals assessed with different sequencing platforms. Linear regression analyses were performed on the genetic effects obtained from the BGISEQ-500 and Blackbird sequencing platforms for maternal metabolites. Similarly, linear regression analyses were conducted on the genetic effect obtained from the Illumina and Ion Torrent sequencing platforms for neonate metabolites.

## Code and Data Availability

The pipelines for performing human genetic analysis using the NIPT data is available in https://github.com/liusylab/NIPT-human-genetics.

Other software and databases used in this study are publicly available, and the URLs are listed below:

SOAPnuke (v1.5.6): https://github.com/BGI-flexlab/SOAPnuke

BWA-MEM (v0.7.17): https://github.com/lh3/bwa

verifyBamID2 (v1.0.6): https://github.com/Griffan/VerifyBamID

BaseVar (v0.8.0): https://github.com/ShujiaHuang/basevar

GATK (v4.1.8.1): https://github.com/broadgsa/gatk/

SAMtools (v1.9): http://samtools.github.io/

BCFtools (v1.9): https://samtools.github.io/bcftools/bcftools.html

bedtools (v2.27.1-65-gc2af1e7-dirty): https://github.com/arq5x/bedtools2/

STITCH (version 1.0.4): https://github.com/rwdavies/STITCH

GLIMPSE (version 1.1.1): https://github.com/odelaneau/GLIMPSE

PLINK (v1.9): https://www.cog-genomics.org/plink/1.9/

PLINK (v2.00a3.6LM): https://www.cog-genomics.org/plink/2.0/

GATK bundle (hg38): https://console.cloud.google.com/storage/browser/genomics-public-data/resources/broad/hg38/v0

Human genome reference (GRCh38/hg38):ftp://ftp.ncbi.nlm.nih.gov/genomes/all/GCA/000/001/405/GCA_000001405.15_GRCh38/seqs_for_alignment_pipelines.ucsc_ids/GCA_000001405.15_GRCh38_no_alt_analysis_set.fna.gz

The low complexity regions of GRCh38: https://github.com/lh3/varcmp/blob/master/scripts/LCR-hs38.bed.gz

The 1000 Genome Project: https://www.internationalgenome.org/

We used Python (version 3.7.6) and R (version 4.1.1) to analyze data and create plots.

## Acknowledgements

The study was supported by Shenzhen Basic Research Foundation (20220818100717002), Guangdong Basic and Applied Basic Research Foundation (2022B1515120080, 2020A1515110859) and National Natural Science Foundation of China (31900487).

## Author Contributions

Conceptualization, S. Liu, S. Huang and X. Jin; Sample collection & Data curation, S. Liu, S. Huang, X. Qiu, X. Jin and H. Li; Investigation, S. Liu, S. Huang, Y. Liu, Y. Gu, X. Lin and Z. Xu; Methodology, S. Liu, S. Huang, R. Davies, A. Albrechtsen and R. Nielsen; Formal analysis, S. Liu, S. Huang, Y. Liu, Y. Gu, X. Lin, X. Lan, L. Li and Zhe Xu; Visualization, S. Huang and S. Liu; Software, S. Liu, S. Huang, R. Davies and A. Albrechtsen; Validation, S. Huang, H. Liu and S. Cheng; Writing-original draft, S. Liu; Writing-review & editing, S. Liu, S. Huang and H. Zhu and G-B Chen; Project administration, S. Liu and X. Jin; Supervision, S. Liu and X. Jin; Resource, S. Liu, X. Jin, H. Li and X. Qiu.

## Competing interest

The authors declare no competing interest.

## Notes

### Competing Interest Statement

The authors have declared no competing interest.

