## Supplementary Information for "Utilizing Non-Invasive Prenatal Test Sequencing Data Resource for Human Genetic Investigation"

### Supplementary Notes
#### BaseVar algorithm
##### Likelihood Function for a single site

$$L(p) = \prod_{i=1}^{N} P(D_i|\ p) = \prod_{i=1}^{N} \sum_{b}^{A,C,G,T} p(b|p)p(D_i|\ b) \quad (1)$$

where $p(b|p) = p_b$ and the genotype likelihood assuming a haploid model is $p(D_i|\ b) = \{1 - \varepsilon_i\ if\ \ D_i = b\ and\ \ \varepsilon_i/3, D_i \neq\ \ b$. $\varepsilon_i$ corresponds to the GATK corresponds to the GATK-recalibrated error rate converted from the PHRED-scale base quality.

Optimization

We obtain the maximum likelihood estimate $\hat{p} = argmax_p L(p)$ using the EM algorithm with starting value computed by the observed allele frequency:

$$p_b = \frac{\sum_{i=1}^{N}\ D_i = b}{N} \quad (2)$$

In the E step, we compute the posterior probability of allele *b* for individual i at a site *j* as one of the four A/C/G/T bases:

$$P(D_i) = \frac{p(b|p)p(D_i|\ b)}{\sum_{b\prime}^{A,C,G,T} p(b\prime|p)p(D_i|\ b\prime)} \qquad (3)$$

We compute the updated allele frequency *p'* in the M step as

$$p'_b = \frac{\sum_{i=1}^{N} P(b|D_i)}{N} \quad (4)$$

When the change in the maximum likelihood is less than 0.001, we terminate the algorithm.

##### Decision of allelic type and confidence of SNP calling: Likelihood Ratio Test

Formulae (1) – (4) can be used for estimation of allele frequencies of all four nucleotides simultaneously, and may result in tetra-allelic and tri-allelic variant calls. We will use this formulation for SNP calling and for identifying potential tri- and tetra-allelic loci. Denote the likelihood value from the four-allelic model in Equation (1) as $f_4$. We iteratively set the allele frequency of one of the four nucleotides to zero to obtain models of tri-allelic loci. Let $\widehat{f_3}(p_x = 0)$denote the maximum likelihood value when the frequency of allele *x* is constrained to be zero. We then compute a log likelihood ratio statistic as:

$$LRT_{4vs3} = -2log\ (\frac{\widehat{f_3}(p_x = 0)}{\widehat{f_4}})\ (5)$$

The tri-allelic model is nested within the tetra-allelic model and, therefore, the distribution of the $LRT_{4vs3}$ statistic asymptotically follows a chi-square distribution with one degree of freedom, under the assumption of a tri-allelic locus. If the p-values of one of the four $LRT_{4vs3}$ test are significant (<$10^{-6}$), the variant will be classified as a tetra-allelic loci. If not, we move on to the test a model of a tri-allelic locus versus a bi-allelic locus, where x if the $\widehat{f_3}(p_x = 0)$ is the allele with minimum likelihood (which results in maximum p-value out of $LRT_{4vs3}$ ) was set as the alternative-hypothesis and the reduced hypothesis is $\widehat{f_2}(p_x = 0, p_y = 0)$ where $p_y$is the allele frequency for allele y.

$$LRT_{3vs2} = -2log\ (\frac{\widehat{f_2}(p_x=0,p_y=0)}{\widehat{f_3}(p_x=0)})\ (6)$$

Again, the distribution of $LRT_{3vs2}$ asymptotically follows a chi-squared distribution with one degree of freedom under the hypothesis of a bi-allelic locus. If the maximum p-value out of the three $LRT_{3vs2}$ is significant, the variant will be classified as a tri-allelic variant. Otherwise, we continue to test the bi-allelic versus mono-allelic assumption, as defined in the equation below, with *y* being the allele with the highest p-value

$$LRT_{2vs1} = -2log\ (\frac{\widehat{f_1}(p_x=0,p_y=0,p_z=0)}{\widehat{f_2}(p_x=0,p_y=0)})\ (7)$$

Formula (8) is also used to quantify the confidence of the SNP call. We keep variants with p-values less than $10^{-6}$.

Note that our method identifies multi-allelic variants. However, since we don't have sufficient validation of the performance for such variants, we focus on reporting results for bi-allelic loci.

#### Power analysis for genome-wide association studies with NIPT data

Power for genome-wide association test using the NIPT data depends on three parameters: effect sample size, minor allele frequency and the phenotypic variance explained by the variant. The derivation of formulation was previously provided by Visscher et al (PMID:28686856) and power is defined as a function of a non-centrality parameter (NCP) associated with the test statistic used to examine whether the genetic effect equals zero. For NIPT data, the effect sample size is a product of the experimental sample size and the imputation accuracy.

$$NCP = n \times r^2 \times q^2/(1 - r^2 \times q^2)$$

$$q^2 = 2 \times MAF \times (1 - MAF) \times \beta^2$$

$$NCP = n \times r^2 \times 2 \times MAF \times (1 - MAF) \times \beta^2$$

$$NCP = n \times R_{imp}^2 \times 2 \times MAF \times (1 - MAF) \times \beta^2$$

$n$: *experimental sample size*

$r^2$: *squared LD correlation*

$q^2$: *proportion of phenotypic variance explained by a causal variant in the population*

$R_{imp}^2$: *saured correlation between the actual and imputed genotypes*

#### Supplementary Figures

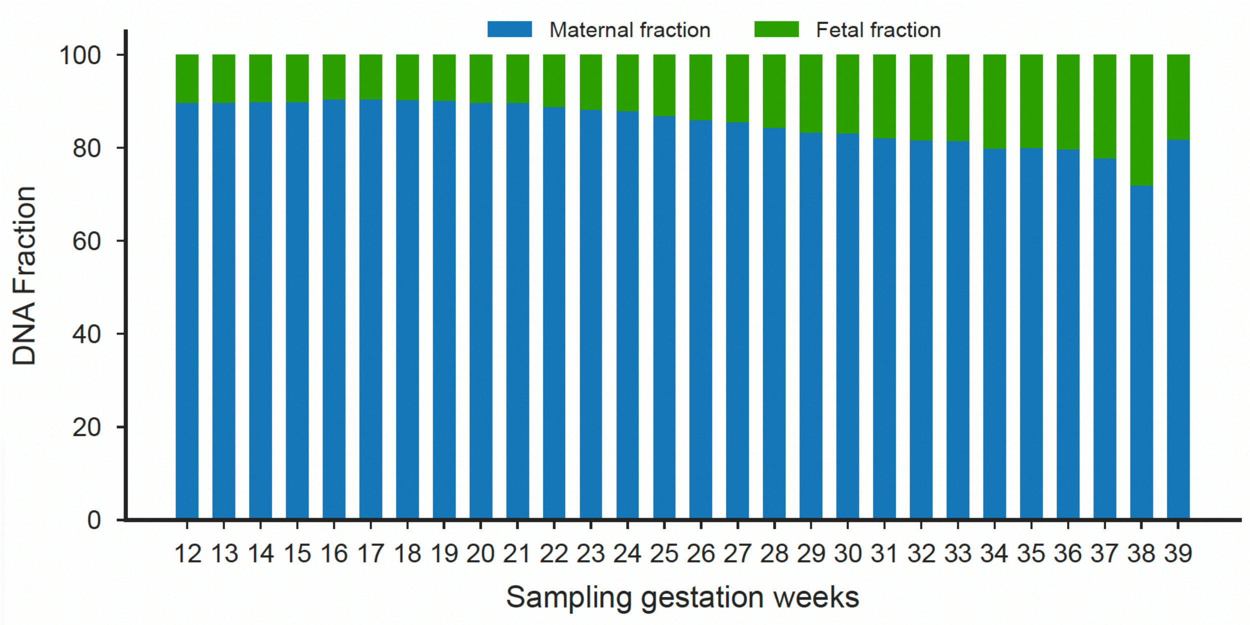

**Figure S1. Distribution of maternal and fetal DNA fraction in NIPT data as a function of sampling gestation weeks**
Statistics are estimated from Y chromosome versus X chromosome coverage from 16,170 pregnancies with male fetus.

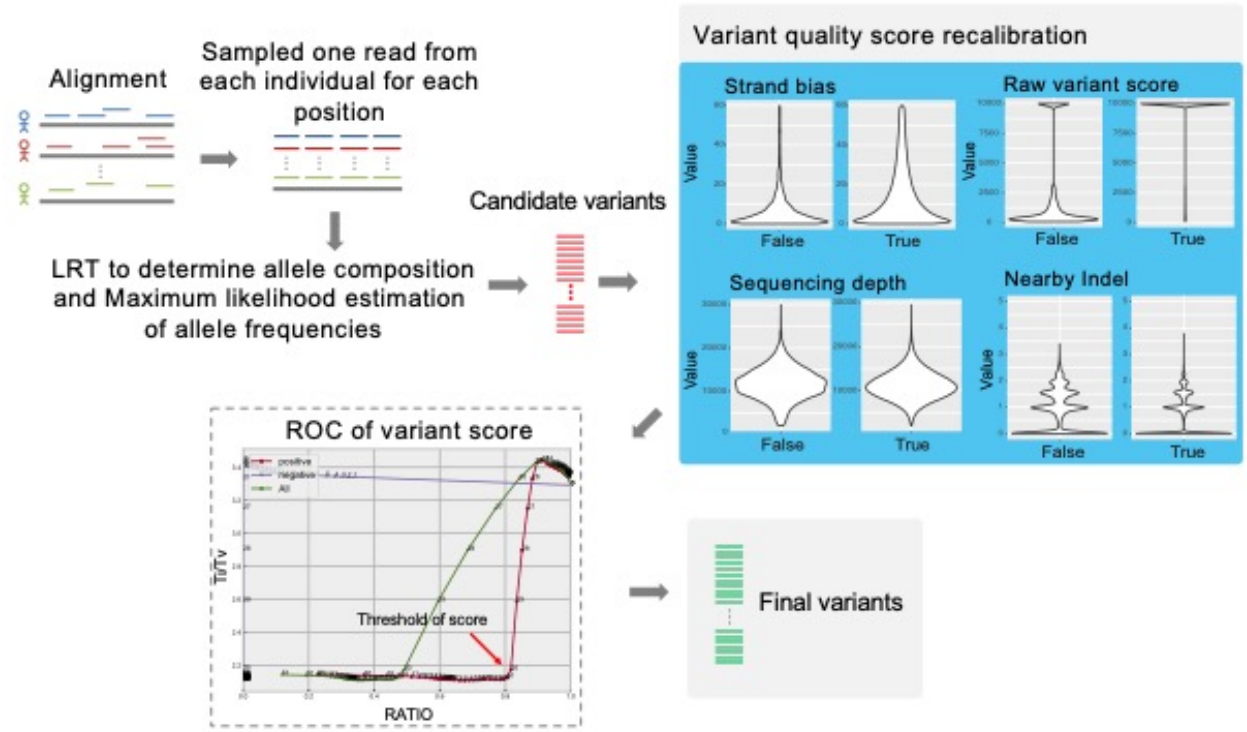

**Figure S2. The BaseVar variation method for variant detection and allele frequency estimation from Non-invasive prenatal test data**

After alignment, one read is sampled from each individual at each position, and an initial allele frequency is estimated by using formula (2). Maximum likelihood estimation of the allele frequency parameters is performed as formula (1) by an EM optimization using formula (3) and (4). Simultaneously, the allele type is determined using multiple rounds of likelihood ratio tests as formulas (5-7). VQSR is eventually applied to determine the best filtration scores for a high-quality set of variants.

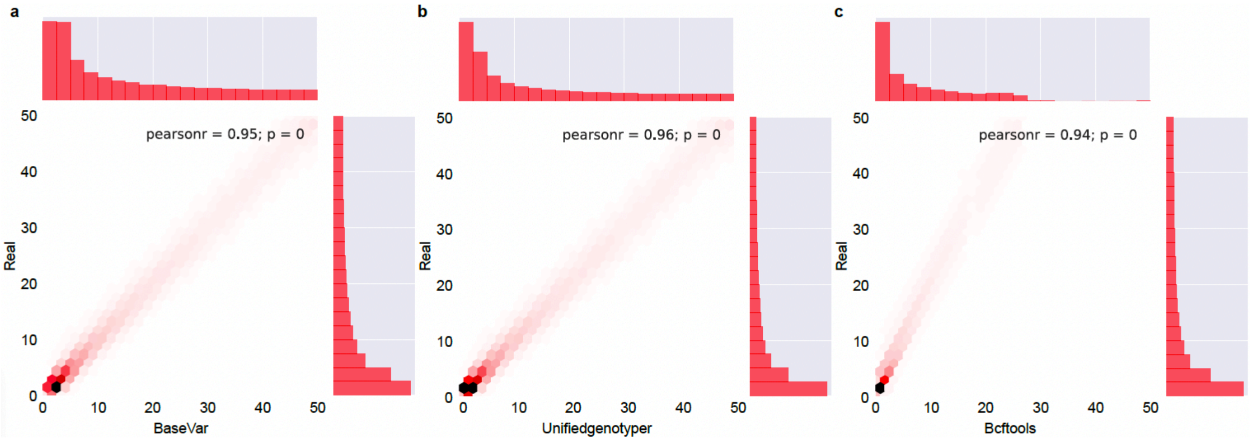

**Figure S3. Consistency of the allele frequency estimation between the true set and the called results by BaseVar, UnifiedGenotyper, and Bcftools**

Reads from 1000 KG Phase II (N=2504, ~7x) was downsampled to 0.1x. Consistency was evaluated by computing the pearson's correlation coefficient between the allele frequency estimates with 0.1x simulated NIPT data and the true allele frequencies reported by 1KGP.

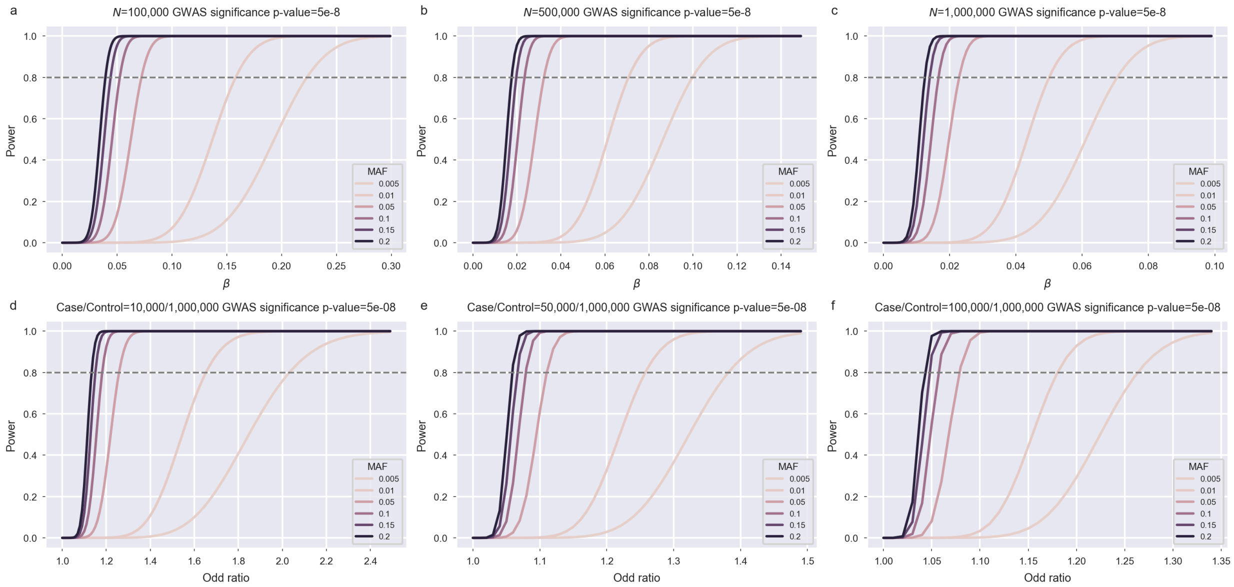

**Figure S4. Power analysis for quantitative and qualitative traits using NIPT data**

(a) – (c)GWAS power estimation for quantitative traits assuming experimental sample size of 100,000, 500,000 and 1 million.

(d) - (f) GWAS power estimation for disease qualitative traits with prevalence rate of 1%, 5% and 10% and an experimental sample size of a million.

The power analysis was conducted using formulas as described in Supplementary Notes, assuming imputation accuracy of 0.8 and a significance level of $5\times10^{-8}$.

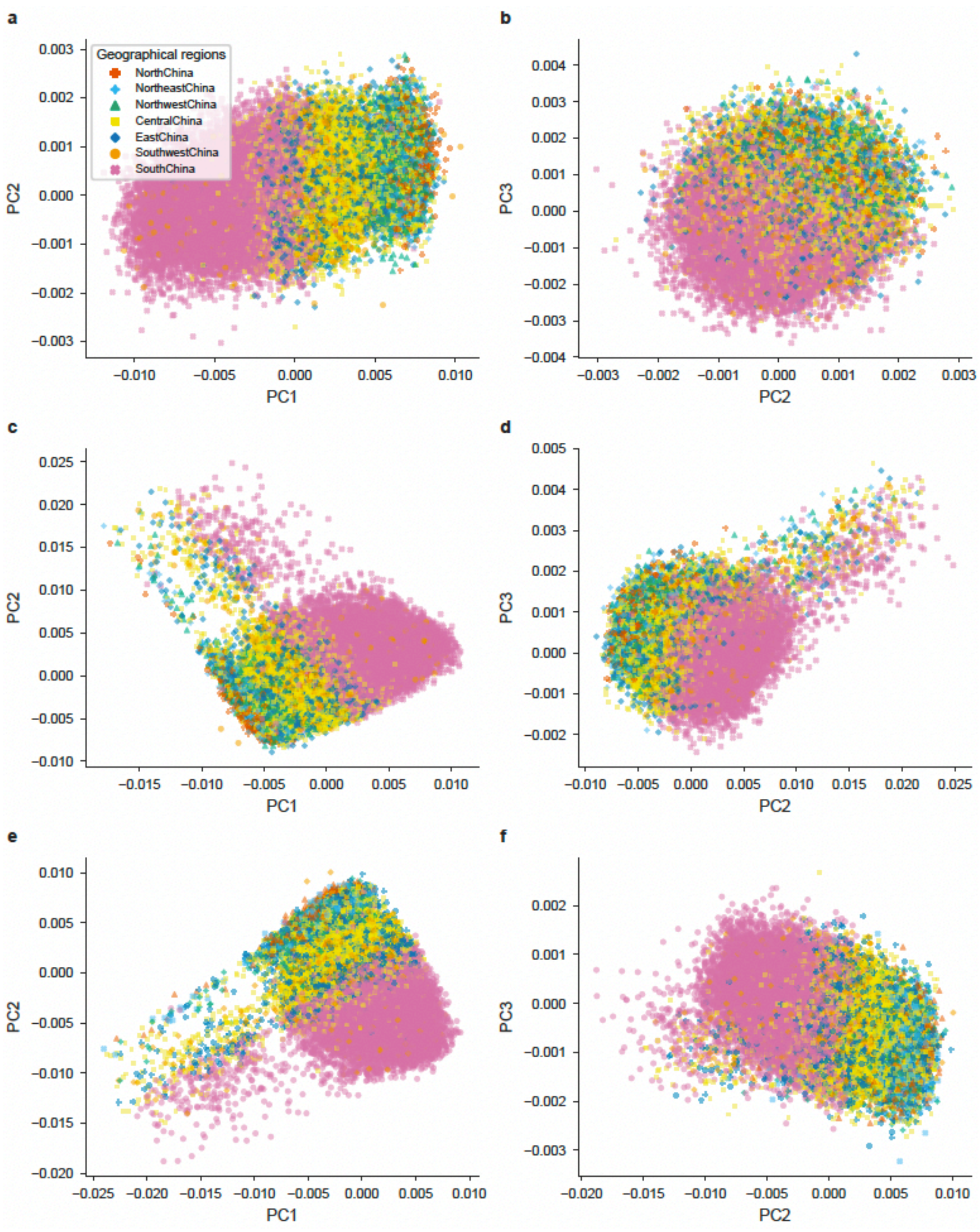

**Figure S5. Principal component analysis using three approaches**

(a) and (b) PCA with PLINK on imputed genotypes

(c) and (d) PCA with PLINK on unimputed genotypes

(e) and (f) PCA with EMU algorithm that did not rely on exact genotypes

#### Supplementary Tables

Table S1. Average sequencing depth for NIPT data provided by three sequencers in China

Table S2. Evaluation of call rate, and allele frequency accuracy as a function of allele frequency on simulation data by sample size of 44K, 140K and 1 million

Table S3. Evaluation of call rate and allele frequency accuracy as a function of allele frequency on simulation data based on allelic type - (bi-allelic, tri-allelic and tetra-allelic)

Table S4. Comparison on computation performance among BaseVar, GATK UnifiedGenotyper and BCFtools

Table S5a. Imputation accuracy for genotype imputation using 1KGP, BIGCS and STROMICS reference panels with QUILT

Table S5b. Imputation accuracy for genotype imputation using 1KGP, BIGCS and STROMICS reference panels with GLIMPSE

Table S6a. Genome-wide association statistics for significant loci with maternal metabolites corresonding to Figure 5a

Table S6b. Genome-wide association statistics for significant loci with maternal metabolites corresonding to Figure 5b
